# Tunable image-mapping optical coherence tomography

**DOI:** 10.1101/2022.10.01.510456

**Authors:** Jaeyul Lee, Xiaoxi Du, Jongchan Park, Qi Cui, Rishyashring R. Iyer, Stephen A. Boppart, Liang Gao

**Author notes:** These authors contributed equally to this work (Jaeyul Lee, Xiaoxi Du).

## Abstract

We present tunable image-mapping optical coherence tomography (TIM-OCT), which can provide optimized imaging performance for a given application by using a programmable phase-only spatial light modulator in a low-coherence full-field spectral-domain interferometer. The resultant system can provide either a high lateral resolution or a high axial resolution in a snapshot without moving parts. Alternatively, the system can achieve a high resolution along all dimensions through a multiple-shot acquisition. We evaluated TIM-OCT in imaging both standard targets and biological samples. Additionally, we demonstrated the integration of TIM-OCT with computational adaptive optics in correcting sample-induced optical aberrations.

---

Optical coherence tomography (OCT), a noninvasive three-dimensional (3D) imaging tool, has been widely used in both basic and translational biomedical sciences [1-4]. Despite significant advances, to acquire 3D images, most current OCT devices require extensive scanning, either in the spatial domain or the spectral domain [5, 6]. The scanning mechanism introduces a trade-off between the imaging speed and image signal-to-noise ratio (SNR), which is particularly problematic for dynamic imaging where the motion of the object can easily blur the image [7-11].

To alleviate this trade-off and enable high-speed 3D microscopic imaging, we previously demonstrated snapshot full-field spectral-domain OCT based on image mapping spectrometer (IMS) [12-14]. By slicing a two-dimensional OCT interferogram in the spatial domain using a custom multifaceted micromirror array followed by dispersing the line images as a spectrum, our proof-of-concept device can capture a 200 μm × 200 μm × 10 μm volume at a rate up to 5 Hz. However, built on a fixed optical architecture, this device suffers from a low spectral resolution—given 480 spatial samplings, the number of spectral bins is limited to 40, leading to a ∼ 5 nm spectral resolution and only a 10 μm depth range in the OCT image. The shallow imaging depth restricts the system from imaging biological tissues. Moreover, the multifaceted micromirror array in the IMS is costly to fabricate, hindering its accessibility to general labs.

To address these technical challenges, herein we present tunable image-mapping OCT (TIM-OCT), which can provide tailored imaging performance for a given application. Rather than using an optically fabricated micromirror array [10-12], we used a phase-only spatial light modulator (SLM) to modulate the incident wavefront in a programmed manner. For example, by displaying a 1D array of linear phase patterns, we can make the SLM act as flat mirrors with varied tilts (Fig. 1a), resembling the micromirror array in the original IMS [15-17]. Alternatively, when displaying a 1D array of linear phase patterns superimposed with quadratic phase patterns, the SLM functions as an array of tilted concave mirrors, focusing the sliced images and redirecting them to varied directions (Fig. 1b). Such a programmable ability allows the IMS to provide a balanced resolution along the spatial, spectral, and temporal dimensions. When combining with OCT, the resultant system can image a 3D volume with either a high lateral resolution or a high axial resolution in a snapshot. The system can also achieve a high resolution along all spatial dimensions through a multi-shot acquisition strategy. We demonstrated TIM-OCT by imaging both standard targets and biological samples.

**Fig. 1.**
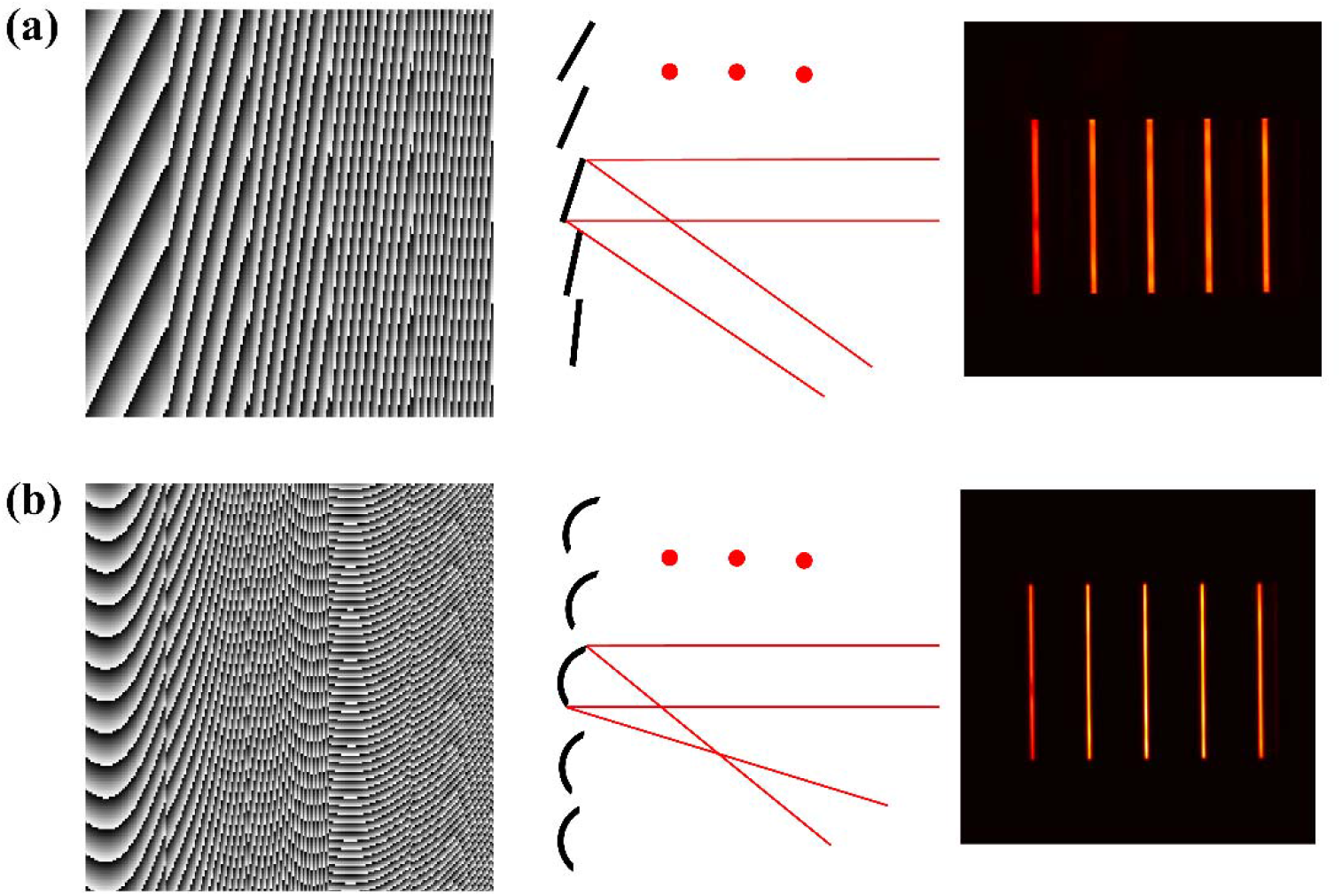
Programmable light guidance using a spatial light modulator (SLM). (a) SLM acts as a 1D flat mirror array with varied tilts, (b) SLM acts as a 1D concave mirror array with varied tilts. From left to right, phase pattern displayed on the SLM, equivalent mirror array configuration, resultant image slices on the camera.

Moreover, we integrated TIM-OCT with computational adaptive optics (CAO) to correct for sample-induced optical aberrations. The application of adaptive optics in OCT has demonstrated immense success in a variety of biomedical applications [18-21]. In particularly, CAO shows unique advantages in data postprocessing flexibility and a reduced system cost [22-25]. However, CAO is extremely sensitive to the phase stability [22, 23], making it challenging for conventional point-scanning OCT systems in dynamic samples. We showed that TIM-OCT can solve this problem by capturing a 3D OCT image in a snapshot, freezing the motion and enabling CAO processing.

The optical setup of TIM-OCT is shown in Fig. 2, and consists of two sub-systems: a full-field spectral-domain OCT system and a tunable IMS. In the OCT sub-system, we use a supercontinuum source (SuperK Fianium FIU-15, NKT Photonics) with a tunable wavelength selection unit as a light source (Fig. 2a). We pass the light through a custom rotating diffuser and illuminate the sample to reduce a speckle noise. The light scattered from the sample is collected by an achromatic doublet lens (focal length, 50 mm) and interfered with the light reflected from the reference arm mirror. The OCT interferogram is then passed to the tunable IMS sub-system. We first filter the light with a polarizer, and then project the image onto a phase-only SLM (HSP1920-488-800, Meadowlark Optics, USA), which has a resolution of 1920 × 1152 pixels and a sensor array size of 17.6 mm × 10.6 mm. We display various phase patterns on the SLM to make it function as an image mapper, analogous to that in the original IMS system. The light reflected from the SLM is then spectrally dispersed by a diffraction grating (GT50-03, ThorLabs, USA; 300 grooves/mm) and imaged by a custom micro-lens array (MLA) (4×8 microlenses; pitch 3 mm; focal length 30 mm), as shown in Fig. 2b. The final dispersed image slices are captured by a CCD camera (hr455MCX, SVS-VISTEK, Germany) as shown in Fig. 2c-2d.

**Fig. 2.**
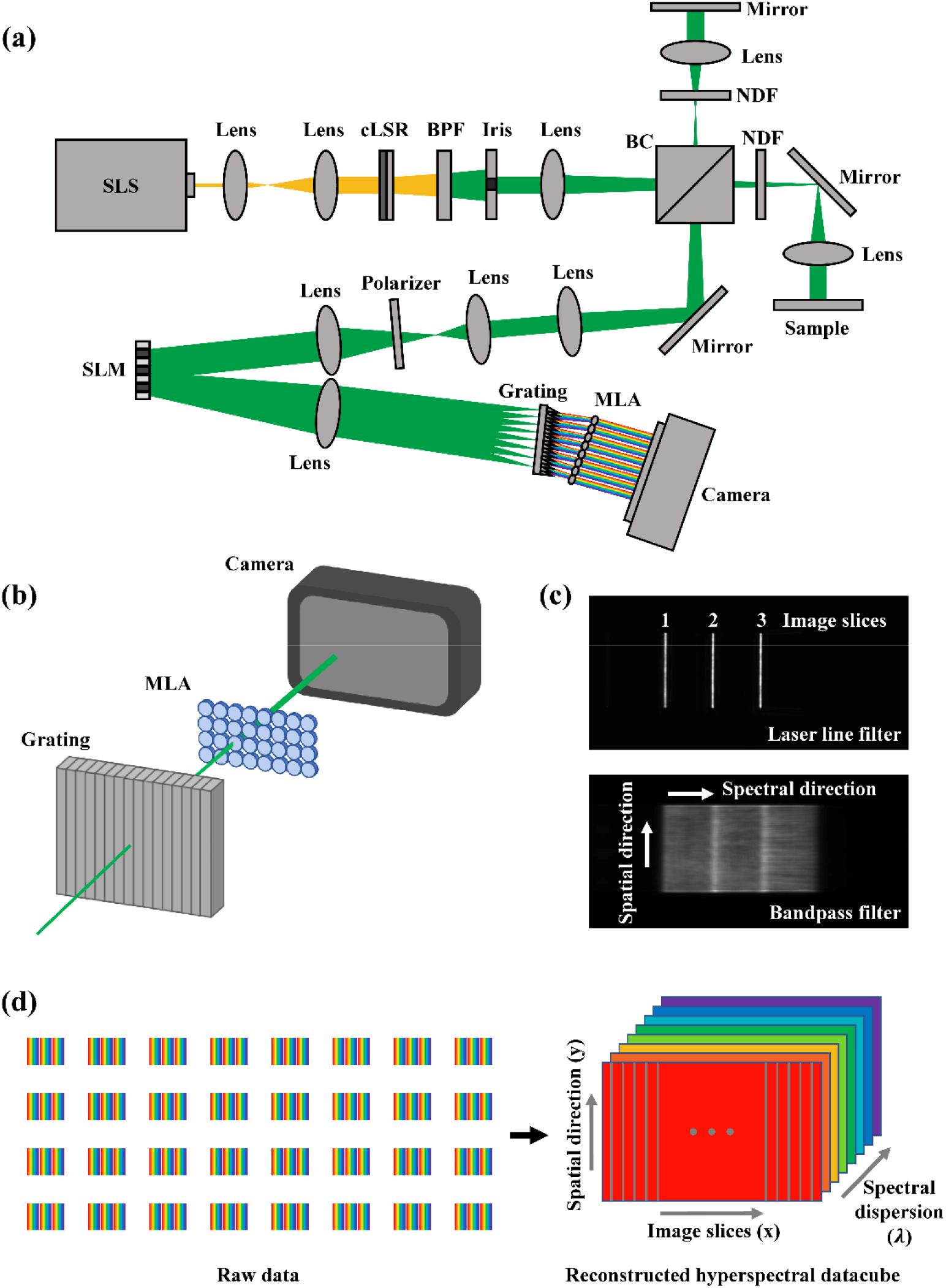
System configuration. (a) Optical setup, (b) 3D view of the grating, MLA, and camera, (c) Representative dispersed image slices under monochromatic (top panel) and broadband illumination (bottom panel, (d) Construction of a hyperspectral datacube. SLS: supercontinuum laser source; AL: aspherical lens; cLSR: customized laser speckle reducer; BPF: bandpass filter; BC: beam combiner; NDF: neutral density filter; SLM: spatial light modulator; MLA: microlens array.

In TIM-OCT, the SLM acts as a mirror facet array, slicing the image into lines and redirecting them towards different microlenses on the array. We operate the SLM in three primary working modes:

<u>Mode 1: Snapshot high-lateral-resolution mode</u>, where the SLM is programed to work as a flat mirror array with varied tilts (Fig. 3a). We divide the SLM into 192 narrow rectangular regions, each having an area of 92 µm (width) × 10.6 mm (length) and displaying a linear phase pattern along a specific direction. We group these SLM regions into six periodic blocks, each block having 32 linear phase patterns with varied orientations, as shown in Fig. 3a-3b. The light reflected from each SLM region is superimposed with the corresponding phase pattern, which essentially directs the light beam towards a microlens on the array. Because phase patterns are repeated across periodic blocks, each microlens on the array collects light from a total of six SLM regions. The spacing between two adjacent reimaged SLM regions is equal to 32 × the width of a SLM region, leading to an effective spectral sampling of 32 after spectral dispersion. Within a single snapshot, the system can capture a spectral datacube (*x,y,λ,*) of dimensions 192 × 155 × 32 (pixel × pixel × effective spectral sampling). To avoid the crosstalk between the adjacent dispersed SLM region images, we filter the illumination using a 10 nm bandwidth filter centered at 550 nm. Given a 270 µm × 170 µm field-of-view (FOV), the resultant OCT image has a lateral resolution of 2.8 µm and an axial resolution of 13.3 µm in air. The volumetric acquisition speed in the snapshot mode is solely dictated by the required exposure time and the camera frame rate (18 frames per second).

**Fig. 3.**
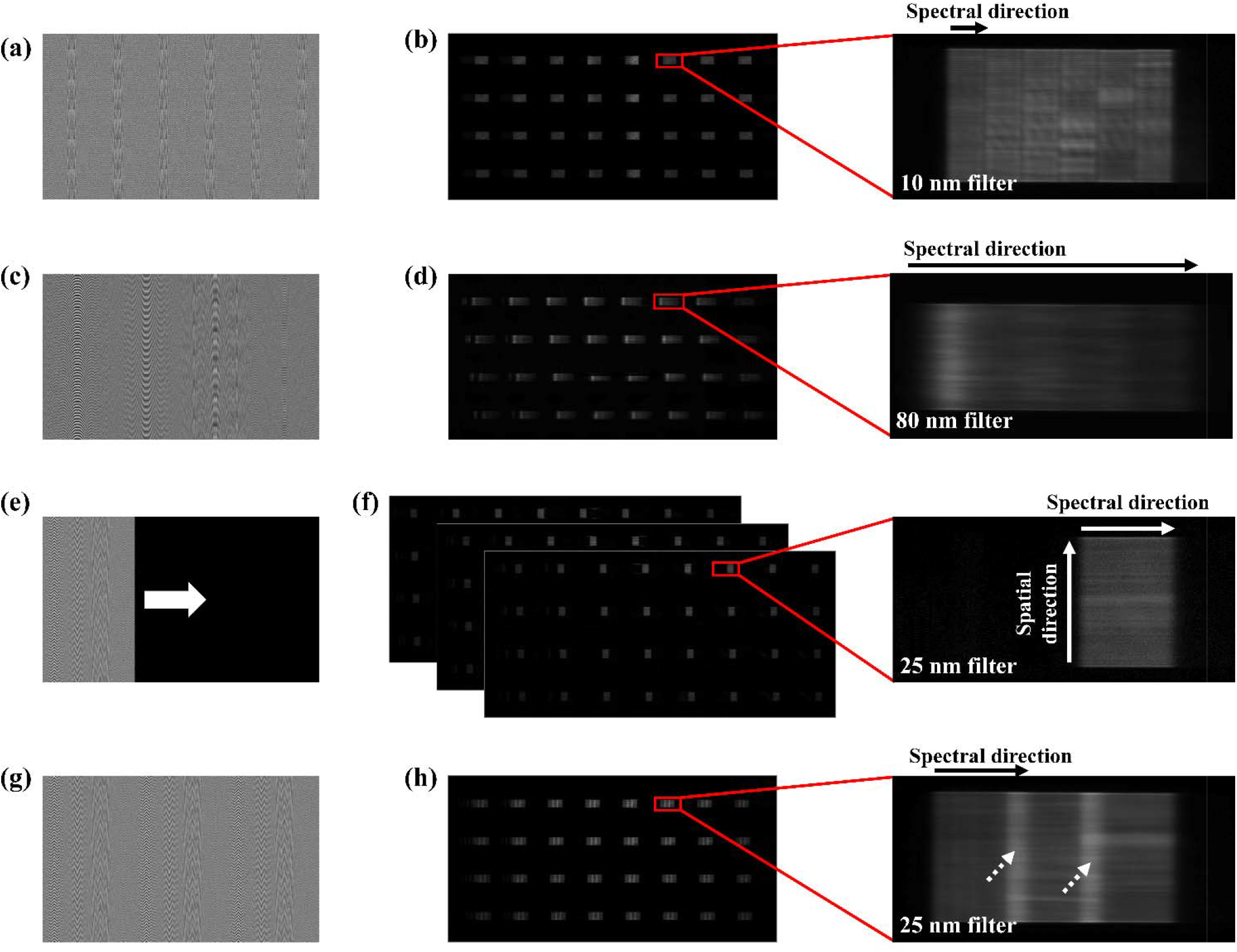
SLM patterns and respective raw camera images in different imaging modes. (a)-(b) Mode 1: Snapshot high-lateral-resolution mode with 192 image slices and a 10 nm spectral bandwidth, (c)-(d) Mode 2: Snapshot high-axial-resolution mode with 32 image slices and a 80 nm spectral bandwidth, (e)-(f) Mode 3: Multi-shot acquisition mode with 96 image slices and a 25 nm spectral bandwidth, (g)-(h) Snapshot acquisition of 96 image slices with spectral crosstalk (arrow-pointed regions).

<u>Mode 2: Snapshot high-axial-resolution mode</u>, where the SLM is programed to work as a concave mirror array with varied tilts (Fig. 3c). We divide the SLM into 32 wide rectangular regions, each having an area of 552 µm (width) × 10.6 mm (length) and displaying a linear phase pattern along a unique orientation and being superimposed with a quadratic phase. The light reflected from a SLM region is first focused into a narrow line (202 µm wide) and then redirected towards a microlens on the array. Because each microlens collects light from only one reimaged SLM region, the corresponding spectrum occupies the entire underlying space, leading to an effective spectral sampling of 160. This allows measurement of a spectral datacube (*x,y, λ*) of dimensions 32 × 155 × 160 (pixel × pixel × effective spectral sampling) in Fig. 3c and 3d. The resultant system can acquire a much broader spectral bandwidth (80 nm) with a 0.46 nm spectral resolution. This combination with the OCT subsystem yields a lateral resolution of 16.8 µm (y-direction) and an axial resolution of 3.4 µm.

<u>Mode 3: Multi-shot acquisition mode</u>, Mode 1 and 2 allow one to tune the spatial resolution along the lateral and axial directions for snapshot acquisition. In Mode 3, one can trade in the temporal resolution for high spatial resolution through multi-shot acquisition. Rather than simultaneously functionalizing all SLM regions, we turn on a subset at a time and capture the resultant spectra in sequence. This increases the spacing between adjacent reimaged SLM regions per acquisition, thereby allowing measurement of a broader spectral range. Figure 3e-3f shows an example of a three-shot acquisition sequence of 96 SLM regions, with each shot capturing 32 line spectra. This mode allows measurement of a spectral datacube (*x,y, λ*) of dimensions 96 × 155 × 54 (pixel × pixel × effective spectral sampling), leading to a 3D resolution of 5.6 µm × 3.4 µm × 10.6 µm in the OCT image. The volumetric imaging speed is 6 frames per second, one-third of the camera frame rate. The illuminating laser power varies by imaging targets from 160 µW (*in-vivo* mouse cornea) to 1.4 mW (*ex-vivo* standard target).

To rebuild a spectral *I*(*x,y,λ*) datacube from the raw measurement, we use a re-mapping algorithm that is described elsewhere [26-28]. Once the *I*(*x,y,λ*) datacube is correctly assembled, we apply a non-uniform discrete Fourier transform (NUDFT) [12] to generate the desired image of 3D sample structure *I*(*x,y,λ*) as illustrated in Fig. 2d. Each location on the sample surface is associated with a separate interferogram, *I*(*x*_*i*_,*y*_*j*_,*λ*) with *i* and *j* corresponding to pixel indices in the *x*- and *y*-dimensions. Conversion of interferograms to A-lines will require re-sampling of raw data from the wavelength *λ* to the wavenumber *k* domain (*k*= 2*π*/*λ*) before Fourier transformation to *z*-space. Repeating the procedure above at each transverse location yields a volumetric image *I*(*x,y,λ*).

To further improve the 3D resolution, we adopt a CAO-based approach to correct for the optical aberrations (Fig. 4). Based on Fourier optics principles, the complex generalized pupil function is proportional to the scaled optical transfer function, which is related to the point-spread-function (PSF) through the Fourier transform. Aberrations thus can be described as a phase term inside the generalized pupil function in a single-pass system. Given a unit magnification in a coherent imaging system, the complex *en-face* wave field in the image space, *Ũ*_*img*_ (*x,y*), is a convolution of the wave field in the object space, *Ũ*_*Obj*_ (*x,y*), with the aberrated complex PSF *P*(*x,y*):

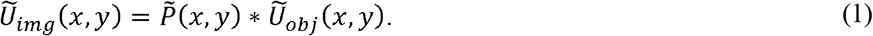

Transforming Eq. 1 to the frequency domain gives:

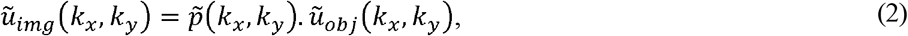

where *k*_*X*_ = 2*π*/*x, k*_*Y*_ = 2*π*/*y*, and 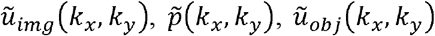 are the Fourier transforms of 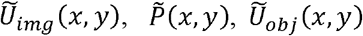, respectively. The coherent optical transfer function 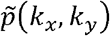can be described by:

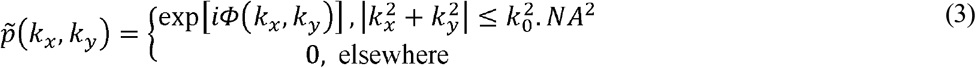

Aberrations of the sample will change only the phase term *Φ*(*k*). Therefore, its effect on the image can be completely reversed by multiplying *ũ*_*img*_(*k*_*X*_,*k*_*Y*_) with the complex conjugate of 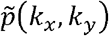.

**Fig. 4.**
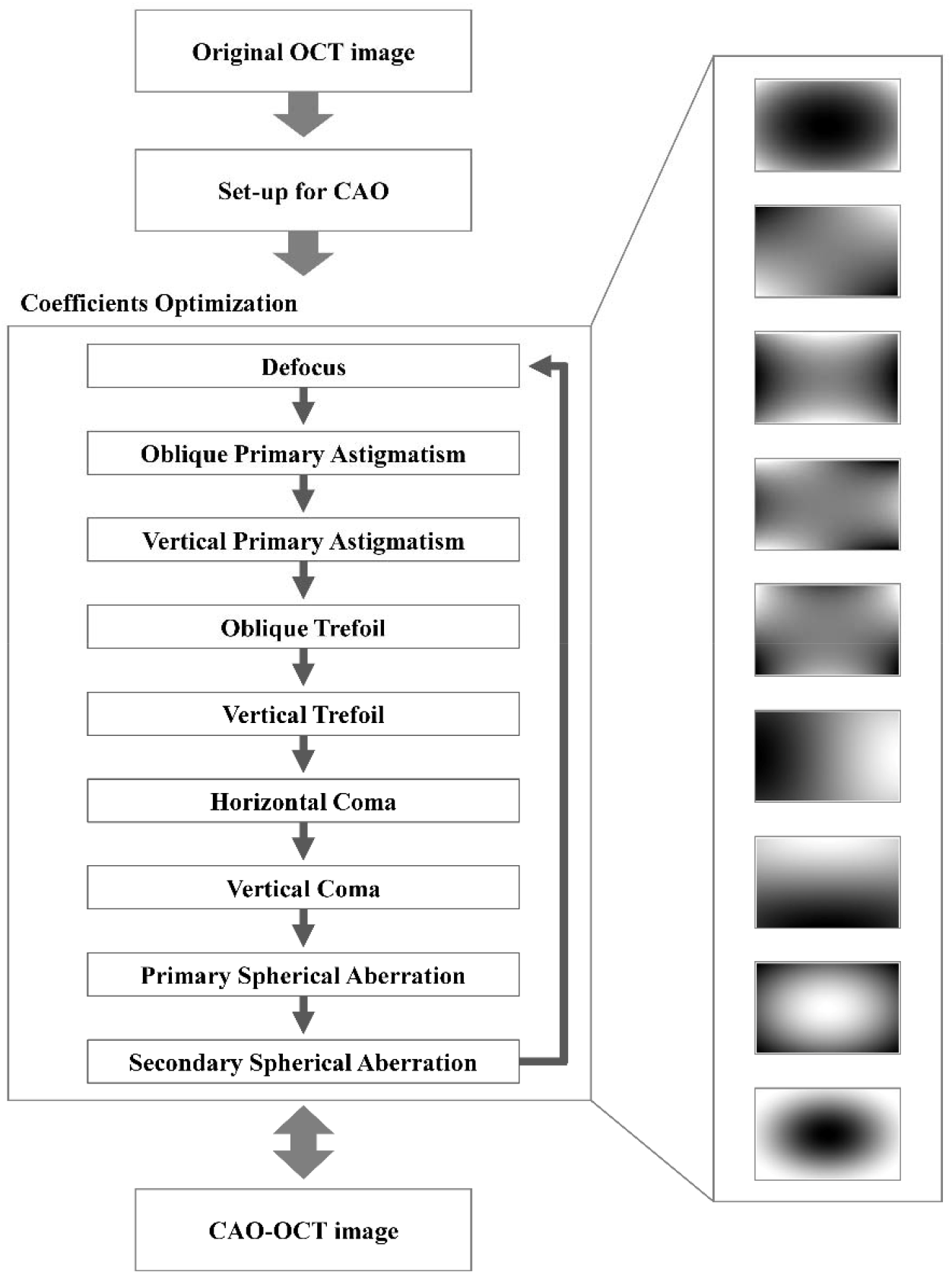
Algorithm of computation adaptive optics (CAO) processing based on the coefficient optimization of Zernike polynomials for aberration-free imaging. The coefficients are modified to optimize the image intensity and sharpness of the OCT image.

To estimate the phase term *Φ*(*k*), we use a CAO-OCT algorithm adapted from Refs [23, 24]. In brief, we first extract a feature-rich *en-face* layer, *Ũ*(*x,y,z*_0_), from a 3D complex image, *Ũ*(*x,y,z*). Fourier transforming the aberrated image *Ũ*(*x,y,z*_0_) yields the aperture function 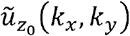. We then expand this pupil function into Zernike polynomials. We follow the pipeline in Fig. 4 and sequentially optimize the phase mask by adjusting Zernike polynomial coefficients from the primary defocus to the secondary spherical aberrations to maximize the sharpness of the resultant image.

To demonstrate the system’s performance in Mode 1 and 2, we imaged a 1951 negative USAF target in air. Figure 5a-5b show the *en-face* and cross-sectional OCT images in the high-lateral-resolution imaging mode, where the *en-face* image presents a high quality image while the cross-sectional image is pixelated due to a lack of spectral sampling. The lateral and axial resolutions are measured to be 2.8 µm and 13.3 µm, respectively. In contrast, when the system is operated in the high-axial resolution mode, it yields a sharp cross-sectional image (Fig. 5c) but a reduced-quality *en-face* image (Fig. 5d). The resultant lateral and axial resolutions are 16.8 µm and 3.4 µm, respectively. It is worth noting that we can switch between these two snapshot working modes simply by changing the phase patterns on the SLM without moving parts.

**Fig. 5.**
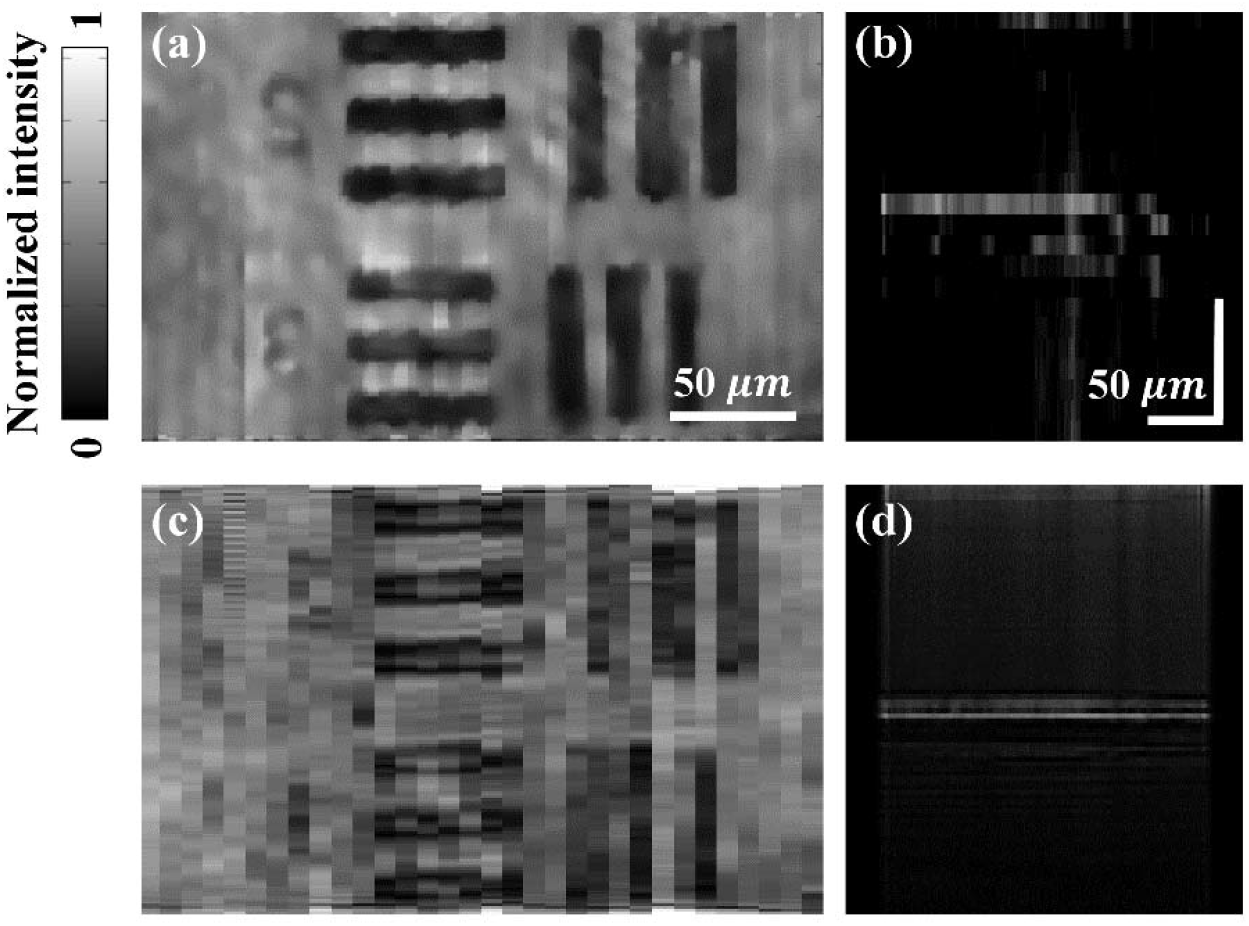
TIM-OCT of a negative USAF target with tunable resolutions. (a) *En-face* image and (b) cross-sectional image in the high-lateral-resolution mode. (c) *En-face* image and (d) cross-sectional image in the high-axis-resolution mode.

We further tested TIM-OCT by performing *in-vivo* imaging of mouse cornea structures in the snapshot *vs*. multi-shot mode. The mouse experiment in this study was conducted in accordance with the guidelines of the UCLA Institutional Animal Care and Use Committee (No. ARC-2021-130) under an approved protocol. A Balb/c mouse (8 weeks old, female) was anesthetized using 0.8% isoflurane delivered at 1 L/min via a snout covering nozzle. The mouse was placed on an electronic heating pad to maintain body temperature, and the mouse condition was constantly monitored during the experiment. We first acquired a corneal image in the snapshot high-lateral-resolution mode (Mode 1), which allows only a 10 nm spectral bandwidth. The resultant cross-sectional intensity image is shown in Fig. 6a, where the epithelium, endothelium, and stromal layers are hardly visible due to the low axial resolution. Next, we switched to the multi-shot acquisition mode (Mode 3) by sequentially capturing the dispersed image slices in three periodic blocks. This increased the spectral bandwidth to 25 nm. The resultant cross-sectional intensity image is shown in Fig. 6b, showing an increased visibility of the corneal layers. Finally, we sequentially scanned a “slit” region on the SLM across the FOV, mimicking the conventional push-broom imaging spectrometers. This led to more extensive scanning but allowed for capturing the full spectral range (80 nm). The resultant cross-sectional image of the epithelium, endothelium, and stromal region is shown in Fig. 6c.

**Fig. 6.**
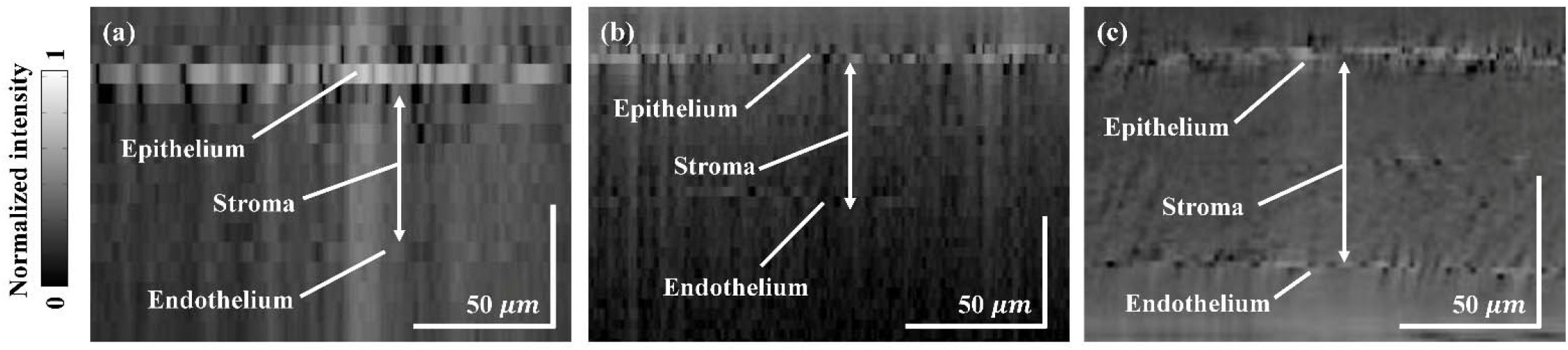
TIM-OCT of an *in-vivo* mouse cornea using snapshot and multi-shot acquisition. (a) Snapshot OCT cross-section image, (b) Three-shot OCT cross-section image, (c) Slit-scan OCT cross-section image.

To demonstrate the integration of snapshot TIM-OCT with CAO, we imaged a negative USAF resolution target with deliberately added aberrations: (1) an in-air 75-µm defocus, (2) a stack of five transparent tape layers on top of the target, and (3) a 1.5 mm thick agarose gel (2.6%) on top of the target. The *en-face* OCT images before/after CAO processing with the applied pupil phase mask are shown in Fig. 7. Figure 7a-7c describes the pupil phase function of the CAO-correction with normalized weights of the nine Zernike polynomials (z4, z3, z5, z9, z6, z8, z7, z12, z15) by the respective aberrations. In all cases, CAO leads to a significant improvement in signal intensities (the line plots in Fig. 7d-7f). Also, CAO processing improves the lateral resolution from 7.0 µm (group 6, element 2) to 3.5 µm (group 7, element 2).

**Fig. 7.**
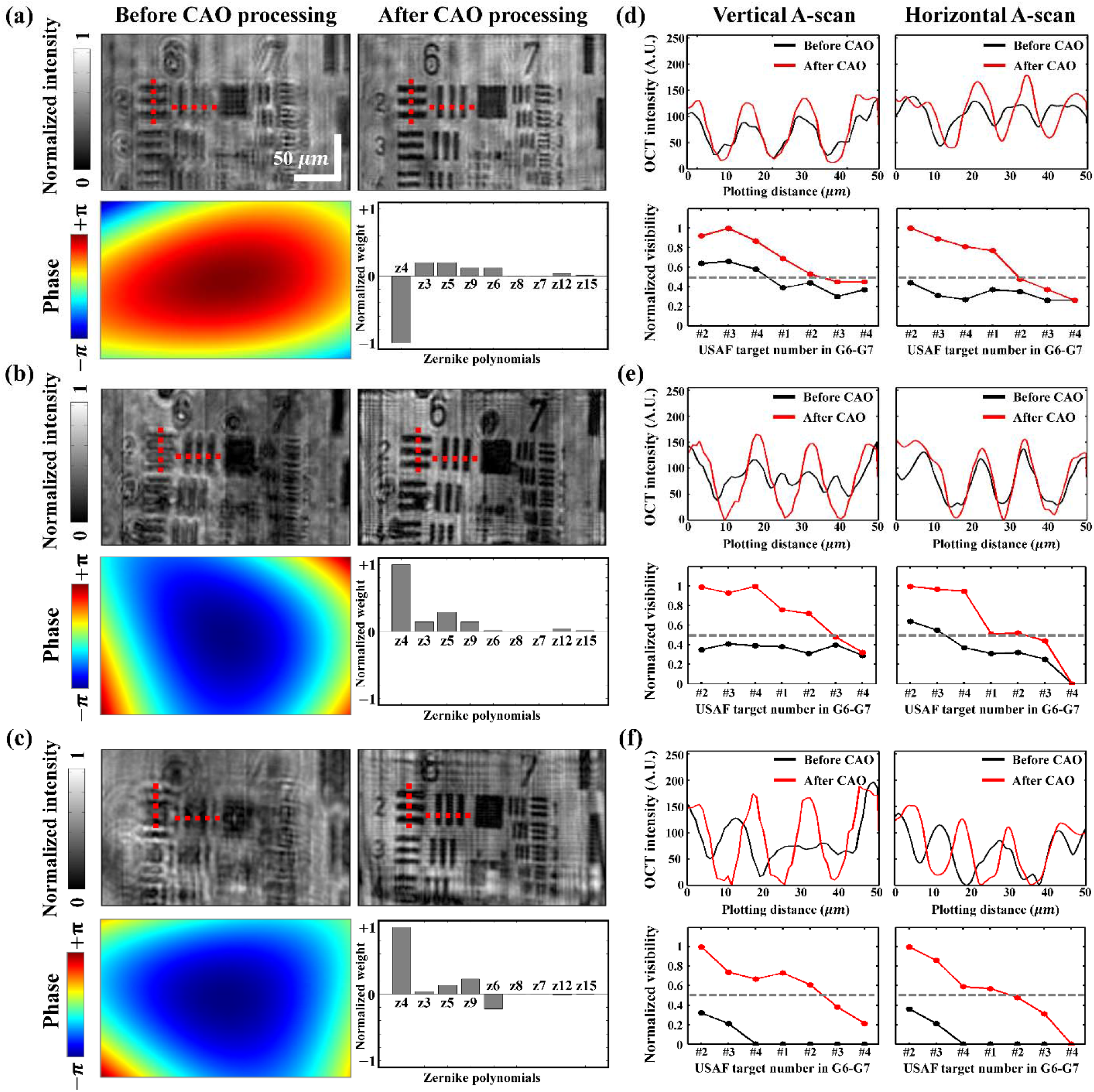
*En-face* OCT images before/after CAO processing and pupil phase function with Zernike polynomials weights under (a) an in-air 75 defocus, (b) a stack of transparent tape strips on top of the target, and (c) a 1.5 mm thick agarose gel (2.6%) on top of the target. (d)-(f) Intensity plots across dashed lines in (a)-(c) and normalized visibility. CAO: computational adaptive optics.

Next, we evaluated the CAO performance with biological samples (Fig. 8). We imaged onion layers and *ex-vivo* chicken breast tissues using TIM-OCT in the snapshot mode, and we optimized the pupil phase function for an out-of-focus depth plane. The CAO corrections were applied to *En-face* images of onion peel and chicken breast tissue at 17.6 µm and 21.1 µm in depth, respectively, with a refractive index of 1.34. The corresponding *en-face* OCT images before/after CAO processing with the applied pupil phase function with normalized weights of Zernike polynomials are shown in Fig. 8. For both biological samples, the image sharpness has been significantly increased after digital correction, as evident by the improved visualization of the onion cellular structures including cell walls and nuclei in Fig. 8a, and chicken tissue structures such as muscle fibers in Fig. 8b. Along the white dashed lines, the sharpness is increased by 2.7 times and 3.3 times, and the intensity is increased by 39% and 32%, respectively.

**Fig. 8.**
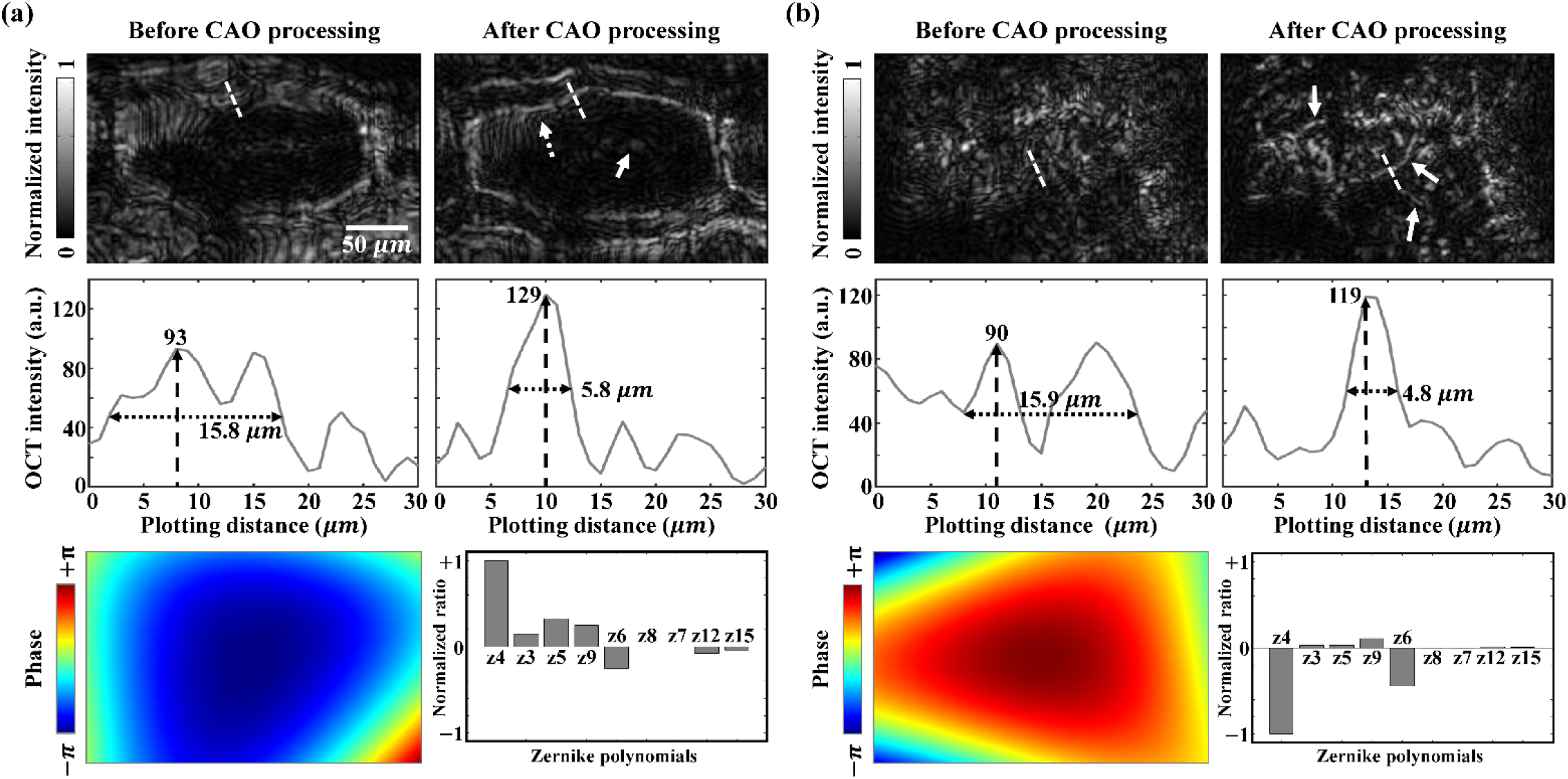
Computational adaptive optics (CAO) processing of TIM-OCT images of biological samples. (a) Onion cells. From top to bottom, *en-face* OCT images before/after CAO processing, with pupil phase function applied with Zernike polynomial weights. The white-dotted arrow indicates the cell wall, the white-lined arrow indicates the cell nucleus. (b) *Ex-vivo* chicken breast. From top to bottom, *en-face* OCT images before/after CAO processing, with pupil phase function applied with Zernike polynomial weights. The white arrows indicate muscle fibers in connective tissues.

In summary, TIM-OCT represents a new category of OCT devices that can be operated in multiple imaging modes, providing tailored imaging performance for target objects. Moreover, the synergy of TIM-OCT with CAO holds great promise in correcting both system- and sample-induced optical aberrations. We expect that TIM-OCT can find broad application in both basic and translational biomedical sciences.

## Data availability

The data that support the plots within this paper and other findings of this study are available from the corresponding author upon reasonable request.

